# Ecology of active viruses and their bacterial hosts in frozen Arctic peat soil revealed with H_2_^18^O stable isotope probing metagenomics

**DOI:** 10.1101/2021.01.25.428156

**Authors:** Gareth Trubl, Jeffrey A. Kimbrel, Jose Liquet-Gonzalez, Erin E. Nuccio, Peter K. Weber, Jennifer Pett-Ridge, Janet K. Jansson, Mark P. Waldrop, Steven J. Blazewicz

## Abstract

Winter carbon loss in northern ecosystems is estimated to be greater than the average growing season carbon uptake. However, most ecosystem carbon measurements neglect winter months since carbon losses (primarily driven by microbial decomposers) are assumed to be negligible at low temperatures. We used stable isotope probing (SIP) targeted metagenomics to reveal the genomic potential of active soil microbial populations under winter conditions, with an emphasis on viruses and virus-host dynamics. Peat soils from the Bonanza Creek LTER site in Alaska were incubated under subzero anoxic conditions with H_2_^18^O for 184 and 370 days. We identified 46 bacterial populations (MAGs; spanning 9 bacterial phyla) and 243 viral populations (vOTUs) that actively took up ^18^O and produced significant CO_2_ throughout the incubation. Active hosts, predicted for 33% of the active vOTUs, were some of the most abundant MAGs and capable of fermentation and organic matter degradation. Approximately three-quarters of the active vOTUs carried auxiliary metabolic genes that spanned five functional categories, including carbon utilization, highlighting the potential impact of viruses in this peat soil’s microbial biogeochemistry. These results illustrate significant bacterial and viral activity and interactions occur in frozen soils, revealing viruses are a major community-structuring agent throughout winter months.

## Introduction

Northern peatlands are important terrestrial ecosystems for carbon (C) storage, estimated to contain one-third of soil C (∼1,000 gigatons) [1], yet the fate of this C is unknown as these soils are experiencing dramatic changes from anthropogenic climate change [2–4]. While soil-warming experiments indicate increased carbon dioxide (CO_2_) and methane emissions under global climate change, it is likely that these C losses from northern peatlands are underestimated because virtually all measurements neglect winter processes [5–8]. Recent estimates of winter C loss alone are estimated to be greater than the average growing season C uptake [9]. While the winter months include large temperature fluctuations and extreme temperature minimums [10], the temperatures found in much of the soil profile of permafrost or seasonally frozen bogs can remain stable and just below the freezing point (□1 to □5°C) [5, 11, 12]. Bacteria have been shown to remain active below the freezing point [13] with both catabolic and anabolic activities observed in frozen soils [14, 15]. Therefore, it is critical to understand the taxonomy, functional capacity, and activities of bacteria and viruses that cause microbial turnover in frozen soils to better predict their biogeochemical implications.

In soils, viruses may play a major role in microbial population dynamics and the fate of soil C, similar to the role they play in marine systems, where viruses kill ∼40% of bacteria daily and sustain up to 55% of bacterial production via C liberation [16, 17]. Soils hold an enormous viral reservoir, but we know remarkably little about their diversity, activity, host interactions, lysis-induced C-cycling, and persistence as compared to other environments [18, 19]. Viruses can affect soil C cycling by stopping microbial metabolism during lytic infections and releasing cell-derived nutrients into the environment [20]. These nutrients can either fuel other organisms’ metabolism or be stabilized via entombing effects [21]. This process can be prolonged by temperate viruses, which remain latent in their host for long periods, before eventually switching into the lytic infection cycle. Viruses can also carry auxiliary metabolic genes (AMGs), host-derived genes that can be expressed during infection and typically aid the infection process by overcoming energetic limitations [22, 23]. While viruses and their impacts have been well-characterized in marine environments via meta-omic approaches [24] and development of tractable virus-host model systems, these linkages remain enigmatic in soil environments. Growing season studies of viruses in northern peatlands indicate they are largely unrelated to other known viruses, highly diverse, endemic to their habitat, infect dominant microbial populations, and carry AMGs that could impact C cycling [25, 26]. The question of whether viruses are active during the 7–9 months of the year when Arctic peatlands are frozen remains unanswered.

Stable isotope probing (SIP) combined with metagenomics is an effective tool for tracking the active microbial taxa in complex communities, linking individuals to specific functions [27–29], or characterizing specific microbes and the viruses that infect them [30]. While many isotope tracing studies use ^13^C-enriched tracers (e.g., ^13^CO_2_, ^13^C-plant biomass, or ^13^C-glucose) to identify specific substrate degraders [28] or assess microbial C use efficiency [31], heavy water (H_2_^18^O) SIP-metagenomics has the unique benefit of labelling all actively growing microbes because water is a universal substrate for nucleic acid synthesis [32–34]. Over adequate time scales, this technique is sensitive enough to label slow-growing or less abundant microbes and identify taxon-specific microbial growth and mortality patterns [35–38], but previously, has not been used to label active viruses.

Here we used heavy-water SIP-metagenomics to label active microbial taxa and viruses and characterize whether viruses play a role in controlling active microbial population dynamics under subzero and anoxic soil conditions. To our knowledge, viral activities have never been confirmed under such conditions, and understanding their impact on microbial activity and soil C transformation over winter months may be pivotal for understanding the future fate of the vast organic C stocks in Arctic peatlands.

## Methods

### Experimental setup

We collected soil samples from the Alaska Peatland Experiment (APEX), part of the Bonanza Creek Long Term Ecological Research (LTER) site southwest of Fairbanks, AK (64.70 °N, □148.3 °W), in a zone of discontinuous permafrost (for overview see Fig. S1). Three peat soil cores were collected from the active layer of a *Sphagnum*-dominated thermokarst bog on June 16, 2013 from the beta site approximately 20 m south of the flux tower. During the week of sampling, midday air temperatures were 27°C, midday surface 10 cm peat temperature was 10°C, active layer depth in the bog was 35 cm, and the water table was 3–8 cm below the surface. The pH was approximately 4.80 (1:1 soil:water ratio). Vegetation and other soil and flux data are given in Waldrop et al., [39]. To collect the cores, moss and peat were removed down to the top of the water table using scissors. The top 20 cm below the water table were collected using a 6.3 cm diameter sharpened steel barrel corer attached to an electric drill. Three cores were collected 1 meter apart along a transect. Cores were stored in mason jars filled with porewater collected from the core hole. Sealed jars were immediately shipped on ice to the U.S. Geological Survey (Menlo Park, CA, USA) where they were homogenized in an anaerobic glovebox maintained at 4°C. Anoxic synthetic porewater was created by freeze-drying filtered (0.45 µm PTFE then 0.2 µm nylon filter) soil porewater collected in tandem with the cores, and then rehydrating the remaining particulates with either heavy water (97 atom% H_2_^18^O, Cambridge Isotope Laboratories) or natural abundance water (control), and then sparging with N_2_ to remove O_2_. Soil subsamples (2 g of soil wet weight) were collected from each core, pressed to remove porewater (using a 5 ml syringe fitted with a nylon screen and a glass fiber filter), and transferred to Wheaton serum vials (10 ml), creating 12 incubation vessels. Vials were sealed with blue butyl rubber stoppers and the headspace was purged (to remove H_2_ from the glovebox atmosphere) by vacuuming and filling with N_2_ (10 inches Hg/5 psi) 10 times in a cold room. Anoxic synthetic ^18^O-enriched porewater was added (2.5 ml) to half of the incubation vials and anoxic synthetic natural abundance porewater was added to the other half using a 5 ml syringe with a 23 G needle that was purged with N_2_. Six parallel samples were set up in a similar manner for headspace gas analysis to quantify CO_2_ production, except these used proportionally larger amounts of soil (18.15 g wet soil) and larger incubation vials (100 ml). All the sample vials were submerged in a glycerol bath at 4°C and the temperature was slowly and steadily reduced to □1.5°C, over 48 h. Samples were then continuously maintained at □1.5°C for 184 d (midyear) and 370 d (full year). At each timepoint, biological replicates (n=3) were destructively harvested and snap frozen in liquid N_2_ and stored at □80°C. Three of the gas production vials were incubated at □20°C as a control.

### CO_2_ production quantification

Gas samples were collected from gas production vials at 10 timepoints over the 370 d incubation. To prevent oxygen from contaminating the incubation vials, a 5 ml syringe with a stopcock and 23 G needle was cleared 3 times with O_2_-free N_2_. The syringe was then inserted into the vial septa and plunged 3 times to mix the headspace; 2 ml headspace was collected, and the stopcock locked. The 2 ml samples were transferred to 10 ml serum bottles that had been purged and then filled with N_2_ (1 atm). Gas samples were analyzed via gas chromatography (SRI 8610GC, SRI instruments, Torrance, CA) to quantify headspace CO_2_.

### DNA extraction and density gradient SIP

DNA was extracted from each replicate using a modified phenol/chloroform protocol [40]. In summary, 0.5 g (+/− 0.01 g) wet soil was added to lysing matrix E tubes, and 100 µl 1x TE (pH 7.5), 150 µl PO_4_ buffer (0.2 M in 1 M NaCl), and 100 µl 10% SDS were added and vortexed. Tubes were bead beat for 30 s at 30 1/s and briefly centrifuged. 0.6 ml phenol:chloroform:isoamyl alcohol (25:24:1) was added, vortexed, and incubated at 65°C for 10 min. Tubes were spun for 5 min at 10K x g, and the supernatant was transferred to a new tube. The bead beat tubes were then re-extracted using 220 µl 1x TE and 80 µl PO_4_ buffers. The supernatant from the first and second extracts were combined in a new 2 ml tube. 550 µl phenol/chloroform/isoamyl alcohol was added, mixed, and centrifuged (10K x g, 5 min). The aqueous top layer was transferred to a new 2 ml tube. 900 µl chloroform:isoamyl alcohol (24:1) was added, mixed, centrifuged (10K x g, 5 min), and the supernatant transferred to a new 2 ml tube. Then, 850 µl chloroform:isoamyl alcohol (24:1) was added, mixed, centrifuged (10K x g, 5 min), and the supernatant transferred to a new 1.7 ml tube. RNAase was added (6.44 µl, 10 mg/ml), mixed, and incubated at 50°C for 10 min. 244 µl 10 M NH_4_^+^ acetate was added, mixed, and incubated at 4°C for 2 h. Tubes were centrifuged at 16.1K x g for 15 min, and the supernatant transferred to a new tube. Isopropanol (670 µl) was added, mixed, and centrifuged (16.1K x g, 20 min). The supernatant was removed, and the DNA pellet dried in a PCR hood for 15 min. 30 µl 1xTE was added, mixed, and the DNA was then stored at □80°C.

Extracted DNA was fractionated via CsCl density gradient ultracentrifugation to separate ^18^O-enriched DNA as described previously [35]. DNA was binned into 5 fractions based on density, and the binned DNA from the two heaviest fractions (medium-heavy [MH; 1.717–1.725 g/ml] and heavy [H; 1.725–1.750 g/ml]) were sequenced.

### Sequencing and metagenome generation

DNA from the SIP fractions was sent to the Keck Sequencing Facility at Yale University. For each sample, 100 ng of DNA was sheared to 500bp using the Covaris E210 (Covaris, Inc., Woburn, MA, USA), followed by a SPRI bead cleanup using Ampure XP (Beckman Coulter Life Sciences, Brea, CA, USA); DNA quality was checked using a Bioanalyzer chip. The sheared gDNA from the 24 samples was then end-repaired, A-tailed, adapters ligated, and sequenced on an Illumina HiSeq 2500 to generate metagenomes (Table S1). Paired-end 151 nt reads were processed in three steps with bbduk v38 (Bushnell, B.): 1) adapter-trimming (ftl=10 ktrim=r k=23 mink=11 hdist=1 tpe tbo minlen=50), 2) PhiX and Illumina adapter/barcode removal (k=31 hdist=1 minlen=50), and 3) quality-trimming (qtrim=r trimq=10 minlen=50). Metagenomes were successfully generated for 23 of the samples (one did not have enough DNA recovered), with 302 Gbp of sequencing data.

### Recovery of vOTUs from metagenomes

Virus-specific informatics were used to increase the number of viral sequences detected in these soil datasets [19]. Processed reads were assembled into contigs using SPAdes v3.11.1 (-- only-assembler --phred-offset 33 --meta -k 25,55,95 –12)[41]. From the 23 SIP-fractionated metagenomes, we assembled 51 487 619 contigs, with 63% of the total reads mapped to the contigs. Contigs were processed with VirSorter (virome decontamination mode) [42] and DeepVirFinder (DVF) [43] to detect viral contigs. We retained contigs that were ≥10kb, sorted into VirSorter categories 1 and 2, and had a DVF score ≥0.9 and p value <0.05. Viral contigs were clustered at 95% average nucleotide identity (ANI) across 85% of the shorter contig [44] using nucmer [45] to generate a nonredundant set of viral populations (vOTUs). vOTU quality was assessed with CheckV (default parameters) [46]. Coverage of the vOTUs was estimated based on post quality-controlled read mapping at ≥90% ANI and covering ≥75% of the contig [44] using Bowtie2 [47]. Coverage was then normalized per gigabase-pair of metagenome and by length of the contig [48]. Activity of vOTUs was determined by a vOTU being present in the H_2_^18^O samples and not present in the paired natural abundance water samples. The vOTUs were annotated using the virus-centric multiPhATE pipeline (default parameters) [49] and the AMG-centric DRAM-v pipeline (with --skip_uniref) [50]. We note that DRAM-v provides AMG scores only for vOTUs detected via VirSorter; AMGs predicted from the 208 vOTUs recovered from DVF were manually curated. For this manual inspection, we removed putative AMGs that were at contig ends or those with annotations from multiple functional categories. To determine the proportion of temperate vOTUs, we searched for genes associated with proviruses, such as integrase and parA [51], leveraged classification from VirSorter (categories 4 or 5) and our genome-similarity host linkages (≤90% similarity), and used two tools — CheckV [46] and PHASTER [52].

### MAG curation and host linking

To generate metagenome assembled genomes (MAGs), read-pairs from the biological replicates were grouped for 8 separate co-assemblies (2 timepoints x 2 SIP fractions x 2 O isotopes) with MEGAHIT v1.1.4 (--k-min 27 --k-max 127 --k-step 10) [53]. Contigs greater than 1 KB were separately binned with Concoct v1.0.0 [54], MaxBin v2.2.6 [55] and MetaBAT v2.12.1 [56]. Genome bins from these three binning tools were refined using the bin_refinement module of MetaWRAP v1.2.1 (-c 50 -x 10) [57] with CheckM v1.0.12 [58]. Only genome bins with at least ‘medium quality’ according to MIMAG standards [59] were retained.

Two methods were used to define MAGs as active: 1) a read-subtraction approach, and 2) a log-fold-change approach. For the read-subtraction approach, contigs from the ^16^O assemblies were used as a reference to subtract the ^18^O reads that aligned with the unlabeled dataset using bbsplit v38 (maxindel=1) [60]. The isotopically labeled reads that did not align with the unlabeled dataset were considered ‘active’ reads and were then processed through the same MAG assembly workflow described above (starting with the MEGAHIT assembly through MetaWRAP refinement); this generated genome bins reconstructed from distinct ^18^O reads from the two SIP fractions at two timepoints.

Refined bins from all 12 groups (8 total + 4 active) were dereplicated into a final set of metagenome assembled genomes (MAGs) using dRep v2.2.3 (-comp 50 -con 10 -p 6 -nc 0.6) [61]. MAG taxonomy was determined using GTDB-tk v0.3.2 [62] with the version r89 GTDB database. Structural annotation was done using Patric [63], and general functional annotation with RASTtk [64] for subsystems within Patric v3 (retaining subsystem variants predicted to be active or likely), and KofamScan v1.1.0 [65] with the KEGG [66] 93 database. Per-sample MAG abundances were determined by aligning each sample’s filtered reads against the MAG genomes using bbmap v38 [67] to obtain total assigned reads per contig and average fold coverage per contig.

For the log-fold-change approach to define active MAGs, we assessed significant (5% false-discovery rate) and positive log2-fold change in ^18^O versus ^16^O read abundances within a time point and SIP fraction. Fold changes were determined using wrench-normalized [68] total assigned reads per MAG with a zero-inflated log-normal model implemented in metagenomeSeq [69].

The vOTUs and MAGs were linked via clustered regularly interspaced short palindromic repeats (CRISPR) spacers and shared genomic content as previously described [70]. Briefly, CRISPR regions were detected in the MAGs using MinCED (options -minNR 2 -spacers) [71] and linked to the vOTUs with blastn (options -max_target_seqs 10000000 -dust no -word_size 7) [72]. In addition, BLAST (options -dust no -perc_identity 70) was used to link vOTUs and MAGs based on shared genomic content [73]. Virus-host abundance estimates were made by summing microbial host abundances at the phylum level in each timepoint and linked vOTU abundances.

### Data availability

The 23 metagenomes were deposited to NCBI under BioProject identifier (ID) PRJNA634918 with BioSample information included in Table S1. Figures were generated with Microsoft Excel and R, using packages Vegan for diversity and pheatmap for heat maps. T-tests and linear regressions were performed using the data analysis package in Excel.

## Results

### Characterization of viruses

To characterize viral activity in Arctic peat soil under winter conditions, we analyzed viral sequences from heavy-water SIP-targeted metagenomic reads. Using two viral detection methods, we identified 5 737 putative viruses (≥5 kb) that clustered into a nonredundant set of 332 vOTUs ≥10 kb (Table S2) that span 66 viral genera (Fig. S2; Table S3). The size range of these vOTUs was 10 039 – 437 858 bp (average 32 954 bp) with 15 vOTUs ≥100 kb, including 5 so-called ‘huge’ viruses (i.e., ≥200 kb) [74]. The vOTUs were well-covered with an average of 17x normalized coverage per metagenome, but with a large range, 3–147x (Fig. 1A). After quality checks, we identified 58 medium–high quality vOTUs, of which four were considered ‘complete’ according to community standards [44]. Genome quality could not be assessed for 93 vOTUs because they did not possess a known viral gene and were detected via a reference-free machine learning method [43]. Annotation of the vOTU genomes with the virus-centric Multiphate pipeline predicted 15 772 genes, of which 61% were novel (Table S4). With the AMG-centric pipeline DRAM-v, we predicted 86 putative AMGs (score 1–3) distilled into five functional categories (carbon utilization, energy, organic nitrogen, transporters, and miscellaneous) from 31 vOTUs (Table S5); 21 of the vOTUs were active and carried 63 AMGs (Fig. 2). To identify temperate viruses, we searched the annotations for genes used in the lysogenic infection cycle and predicted nearly half (43%) of our vOTUs were temperate viruses. More than half (59%) of these temperate viruses were active, and the majority (80%) had at least one member of their population integrated at the time of sampling (provirus; Table S6).

**Figure 1.**
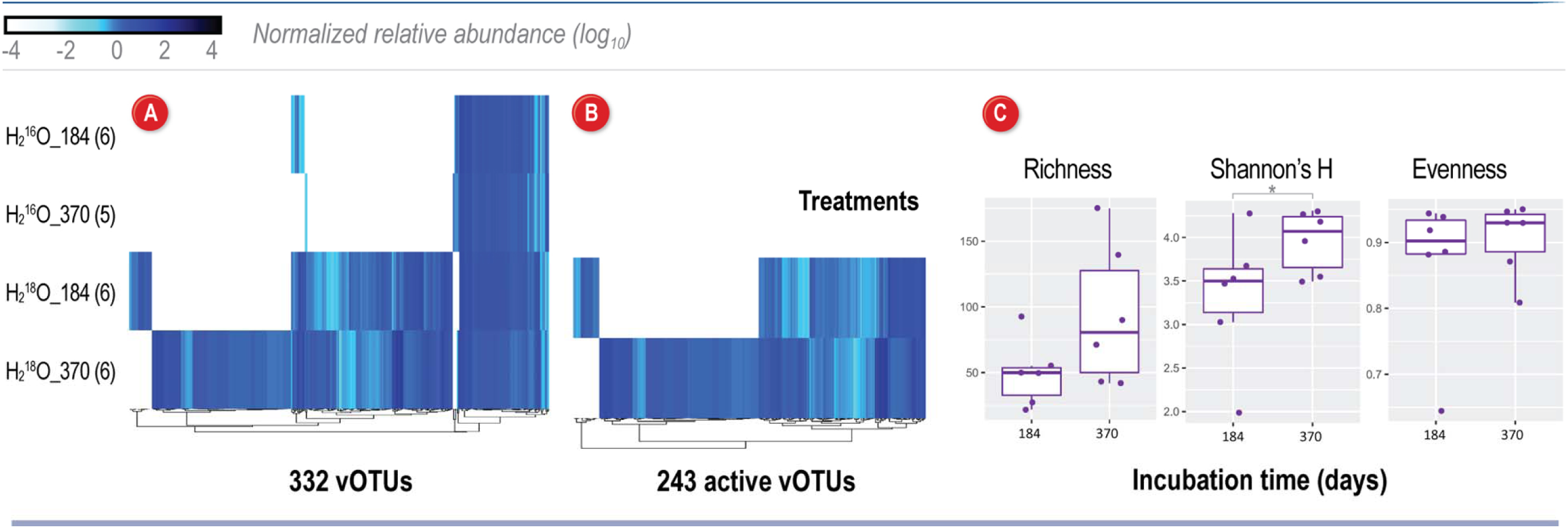
Assessment of viral community structure and activity after a ^18^O water incubation in Arctic peat soils. (A) Viral sequences were identified from 23 samples grouped by two treatments (natural abundance water “H_2_^16^O”, and heavy water “H_2_^18^O”) and two time points: 184 days and 370 days. Number of replicates is indicated in parentheses. Relative abundances of all 332 vOTUs identified in the peat soils, clustered by abundances in each treatment/timepoint. (B) Relative abundances of 243 vOTUs considered ‘active’ due to DNA ^18^O enrichment patterns. Relative abundance for each vOTU (illustrated by blue gradient) was normalized by metagenome size (total base pairs) and contig length, and reads were mapped to the contig if they shared ≥90% average nucleotide identity and had ≥85% alignment fraction. (C) Diversity metrics for the 243 active vOTUs. Box plots show the median, upper and lower quartile range, and the variance among the samples. Asterix denotes significance (p <0.05).

**Figure 2.**
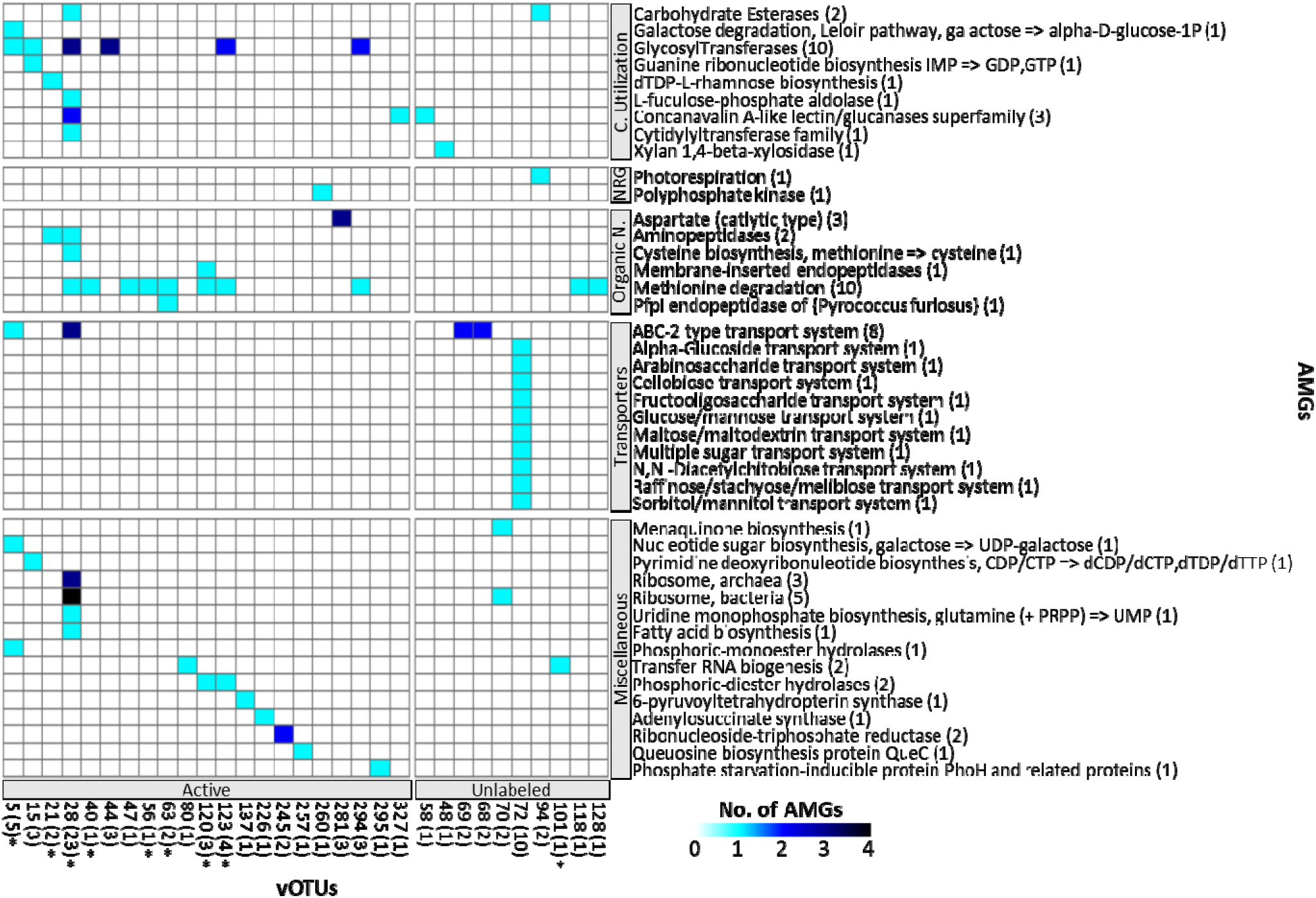
Putative auxiliary metabolic genes associated with peat soil viral genomes. Heatmap of 31 vOTUs carrying confidently predicted auxiliary metabolic genes (AMGs) (Dram-v score 1–3) and their annotation. The vOTUs are grouped by active or unlabeled (see Table S3 for additional detail) with the sum of AMGs per vOTU indicated in parenthesis and those with an asterisk are linked to an active MAG. AMGs are grouped by functional category — carbon utilization, energy generation, organic nitrogen use, transporters, and miscellaneous.

Significantly more vOTUs were observed in the SIP fractions from heavy-water incubations relative to control (natural abundance water) incubations, confirming that many viruses incorporated ^18^O into their DNA and were actively replicating (Fig. S3). A majority (73%) of the vOTUs found in the dense ^18^O SIP fractions were active at some point over the 370-d incubation, with about half active the full year and the other half only active in 184 d or 370 d samples (Fig. 1B). We measured active vOTU diversity to assess changes in the viral community structure from midyear to a full year of incubation. The Shannon’s H metric indicates a significantly (*p* ≤0.05) more diverse viral community at 370 d compared to 184 d (Fig. 1C). Shannon’s H diversity, which includes both richness and evenness, was driven by an average increase of more than 60% for vOTU richness from 184 to 370 d. Of these, 64 vOTUs became more abundant from 184 to 370 d, 110 became newly active, and 18 were no longer detected as labeled at 370 d.

### Host characterization based on SIP-metagenomics

To identify potential viral hosts, we used a suite of metagenome assembly and binning methods which yielded 153 medium to high quality MAGs, spanning 16 bacterial phyla and 1 archaeal phylum (Table S7; GTDB taxonomy). The dominant phyla detected were Chloroflexota, Acidobacteria, and Myxococcota (formally Proteobacteria). Incubation with heavy water indicated 30% (46) of these MAGs were actively growing, spanning 9 bacterial phyla, with the most abundant active populations from the Acidobacteria, Bacteroidetes, and Firmicutes. By sequencing both unlabeled and isotopically labeled samples, we gained insight into genetic repertoire from both active and inactive bacteria (Table S8) but focused our efforts on characterizing the active bacterial lineages and metabolisms that viruses may be altering over winter months. We used CO_2_ production measurements to confirm microbial activity and positive fluxes occurred continuously throughout the □1.5°C incubation, but not from the □20°C control incubation (Fig. 3).

**Figure 3.**
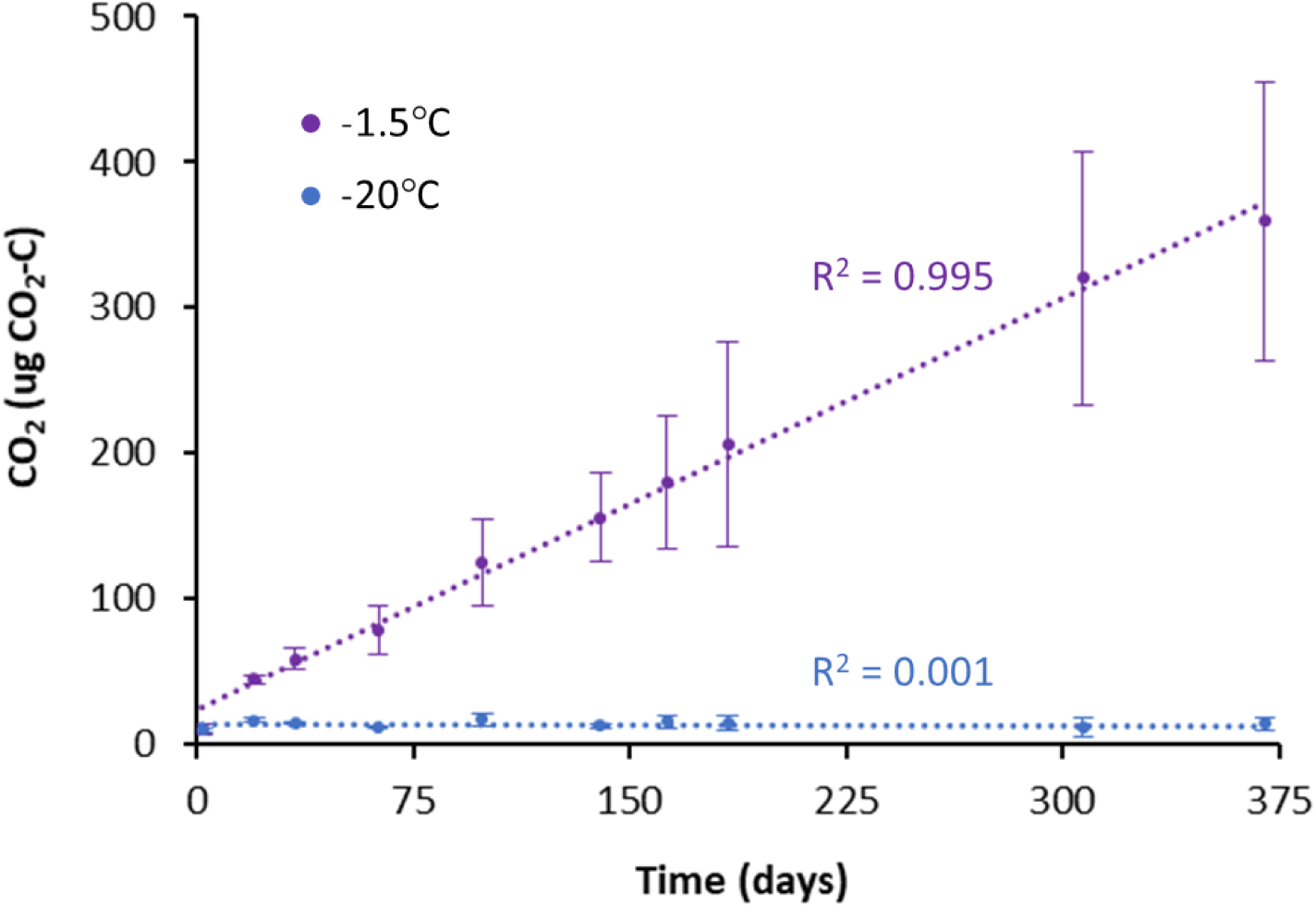
CO_2_ production. Cumulative CO_2_ production in soil incubated at □1.5°C (experimental conditions) and □20°C (control). Error bars show standard error (n=3) and R^2^ is shown for each linear regression.

### Linking vOTUs to MAG hosts

To understand the influence of the viruses on microbial dynamics during the winter season, we predicted potential microbial hosts via two different *in silico* approaches based on sequence similarity. First, we identified 11 879 CRISPR spacers in 136 of the MAGs and used these to link 10 vOTUs to 6 MAGs from 4 bacterial phyla (Fig. 4, Table S9). Leveraging the SIP activity patterns, we found most of these identified linkages connected active vOTUs and active MAGs (12 linkages between 8 active vOTUs and 5 active MAGs, 1 linkage between an unlabeled vOTU and an active MAG, and 3 linkages between unlabeled vOTUs and unlabeled MAGs). In a second approach, we used vOTU-MAG genome sequence similarity and recovered 798 similarity matches that linked 141 vOTUs to 65 MAGs from 10 bacterial phyla (Fig. 4; Table S10). Combined, the two virus-host linkage approaches indicated 318 unique connections between 145 vOTUs and 65 MAGs spanning 10 bacterial phyla. A significantly higher proportion of these vOTU-host matches were between active populations (Table 1). Notably, four vOTUs (#s 153, 161, 162, and 270) had a broad-host range, and were linked to bacteria from several bacterial phyla (three of these vOTUs infected two phyla and one infecting four phyla; Fig. 4). Two of these broad-host-range vOTUs (153 and 270) were active and linked to only active MAGs from bacterial phyla Bacteriodota and Firmicutes.

**Figure 4.**
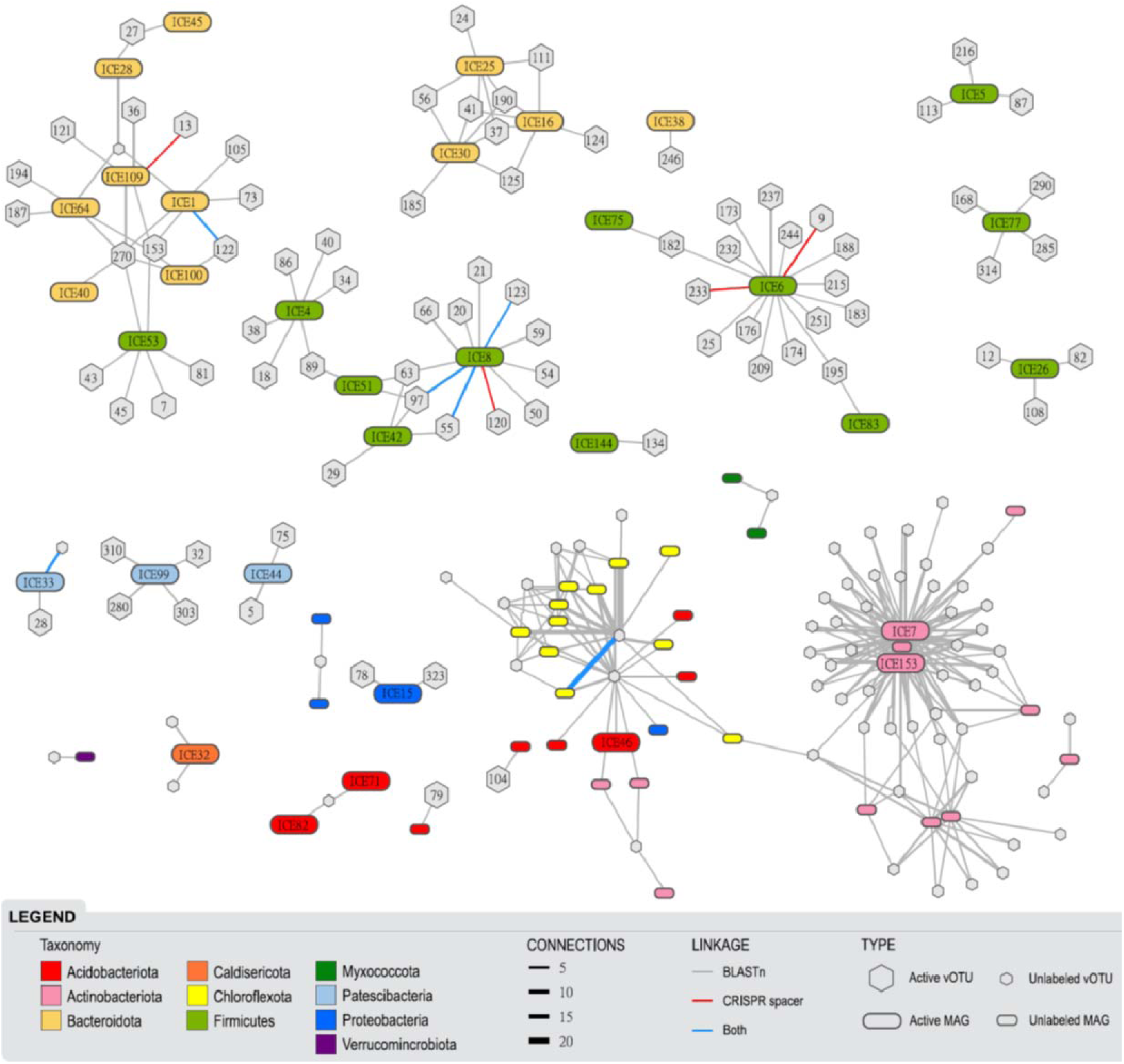
Virus-host linkages in Arctic peat soils incubated with ^18^O enriched water. Network diagram illustrating vOTUs and their predicted bacterial hosts, colored by bacterial phyla. Active versus inactive vOTUs and MAGs (based on ^18^O enrichment) are indicated by large named or small hexagon/rounded rectangles, respectively. Lines represent linkages between a vOTU and a bacterial MAG, thickness denotes the number of connections, and are colored by the identification approach used: similarity in genomic content (gray), CRISPR spacer match (red), or both (blue).

**Table 1.** Linkages between ^18^O labeled and unlabeled viruses and bacteria in a SIP incubation of sub-zero, anoxic Arctic peat soils. Viral operational taxonomic units (vOTUs) were linked to metagenome assembled genomes (MAGs) via nucleotide similarity and using CRISPR spacers. ‘Active’ vOTUs and MAGs were defined based on assimilation of ^18^O enriched water (heavy water) into DNA. In total, there were 814 linkages from active vOTUs to active MAGs (group 1), active vOTUs to unlabeled MAGs (group 2), unlabeled vOTUs to active MAGs (group 3) and unlabeled vOTUs to unlabeled MAGs (group 4).

| Group | Description | No. of vOTU-MAG matches | Unique vOTU-MAG matches | vOTUs | MAGs |
| --- | --- | --- | --- | --- | --- |
| 1 | Active vOTU-active MAG | 164 | 107 | 74 | 37 |
| 2 | Active vOTU-unlabeled MAG | 2 | 2 | 2 | 2 |
| 3 | Unlabeled vOTU-active MAG | 226 | 77 | 44 | 11 |
| 4 | Unlabeled vOTU-unlabeled MAG | 422 | 132 | 58 | 30 |
| **Total** |  | **814** | **318** | **178** | **80** |

All 145 of the vOTUs we identified as host-linked may therefore be classified as dsDNA bacteriophages (since the vOTUs were linked to MAGs with a bacterium taxonomic assignment). These represented the majority (88%) of the host-virus matches, and almost all (92%) of the unlabeled vOTUs that were linked to unlabeled MAGs. In our soil incubations, Actinobacteriota (56%), Chloroflexota (24%), and Firmicutes (12%) were the most ‘infected’ bacterial phyla (i.e., with the most vOTU-MAG linkages). Firmicutes (55%), Bacteroidota (34%), Patescibacteria (9%), and Proteobacteria (2%) were the only phyla that had active MAGs linked to active vOTUs. Of the active populations, 81 (33%) vOTUs and 33 (51%) MAGs were linked, with the top 15% most abundant active vOTUs predicted to infect Firmicutes and Bacteroidota, and the abundances of both these vOTUs and their host populations increased over the year incubation (Fig. 5).

**Figure 5.**
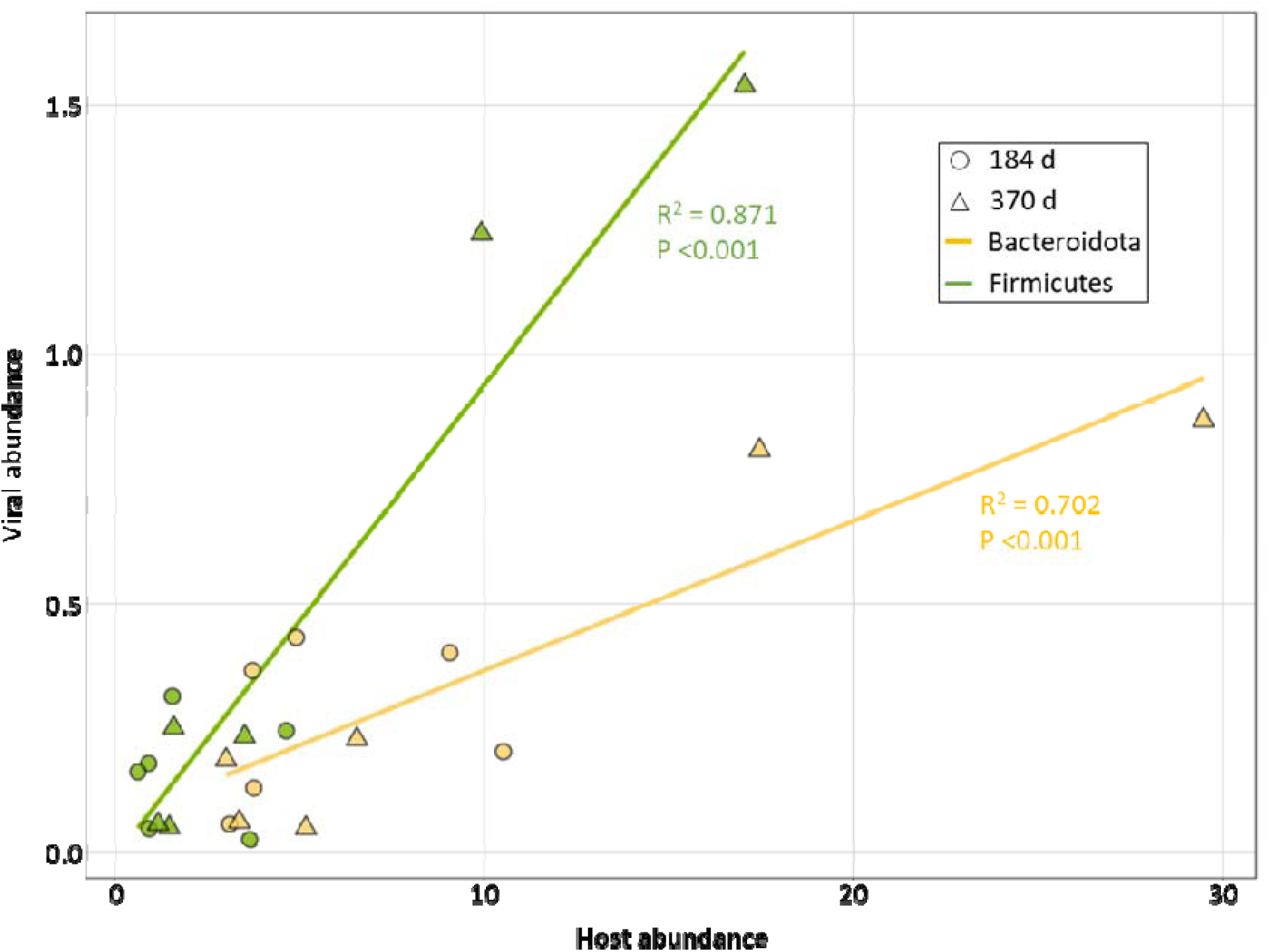
Abundances of active viruses and their predicted bacterial hosts. Average virus:host abundance ratios for bacterial phyla Bacteroidota (yellow) and Firmicutes (green) from heavy-water treatment samples at 184 (n=6) and 370 days (n=6). Host abundance and the abundance of viruses for that host were calculated as the mean coverage depth from metagenomic read mapping, normalized by the number of reads in the sample.

## Discussion

Quantifying and predicting the drivers of C loss in northern latitude peatlands underlain with permafrost is a complex and critical issue, especially as these environments continue to be disproportionately impacted by climate change. Microbes largely govern C release from Arctic soils, and recent work has highlighted the important role viruses could play in microbial C processing and helping their hosts adapt to subzero temperatures [25, 26]. What is not well understood is what proportion of these viruses are active, how the viral community dynamics change over time, and how long viruses persist in frozen anoxic peat soils. To our knowledge, this is the first study to address these concerns, using heavy water SIP-metagenomics to directly identify active microbes and their associated viruses and characterize their dynamics from a half of a year to a full year under subzero and anoxic conditions.

### Viral activity in cryoecosystems

Over the course of a year, we found hundreds of active vOTUs, in stark contrast to the general paradigm that frozen soils have little to no activity [75]. A third of these active vOTUs were linked to active MAGs, highlighting that not only are microbes active in these anoxic subzero temperature soils, but their viruses are also active and likely impacting soil microbial community assembly and ecosystem biogeochemistry, as previously proposed [25, 26, 76, 77]. We observed an increase in vOTU richness and abundance from midyear to a full year of incubation, although vOTU evenness remained the same, likely due to a combination of proviruses replicating with active microbes and many generations of virions actively infecting newly active host(s). The few vOTUs that decreased in abundance or were present midyear and not detected after a full year, could have been isotopically enriched then some persisted in the soil as environmental DNA (eDNA), or as inert virions from internal nucleic acid errors (e.g., lethal mutations) or structural damage, or as competent virions that were no longer infective due to their host evolving resistance (e.g., the ‘host-virus arms race’ that has been observed in other systems)[78].

While most viruses are host-specific, having a broad host range can influence a virus’ ecological significance. Generalist viruses that can infect more than one species of host are thought to have different abundance patterns, infection efficiency, movement within an ecosystem, and other unique attributes [79]. We identified fifteen generalists and four highly promiscuous generalist vOTUs that were predicted to infect hosts from different phyla. In addition, these four generalists were some of the most abundant vOTUs, suggesting that having a very broad host range may offer an advantage in these subzero and anoxic soils. Recently, Malki et al. [80] showed that four viruses from an oligotrophic lake in Michigan could infect bacteria across several phyla including Proteobacteria, Actinobacteria, and Bacteroidetes. Interestingly, the host phyla identified by Malki et al. are the same bacterial phyla that our generalists vOTUs infected, with the addition of Firmicutes and Chloroflexota (also predicted by our network analyses, see Table S11; Fig. S4).

### Putative viral influence on winter biogeochemistry

Soil viruses shape the abundance, diversity, and metabolic outputs of their microbial hosts. As recent literature has made clear, viruses may modulate ecosystem biogeochemical processes via an intriguing mechanism—reprogramming host metabolism with AMGs during infection [24]. The metabolic functions and ubiquity of AMGs can vary by environment, with marine viruses carrying AMGs for photosynthesis, central carbon metabolism, sulfur cycling, and nitrogen cycling [24]. Soil viruses, although understudied, appear to carry AMGs for polysaccharide degradation and sporulation [25, 26, 81]. In our cryoecosystem, we identified carbon degradation AMGs (e.g., galactose degradation and Xylan 1,4-beta-xylosidase) that could provide fitness advantages for utilizing much-needed carbon sources, as well as genes for scavenging (e.g., ABC-2 type transport system) that may provide their host an avenue to acquire essential nutrients [82]. Additionally, AMGs such as methionine degradation [83] and Glycosyl transferases [84] were found in high abundance which can help during infection.

To better understand the roles viruses play in controlling central biogeochemical processes in Arctic peat under winter conditions, we assessed the active bacterial phyla that were virus-infected. Firmicutes had the largest increase in abundance from midyear to a full year and had the most viral infections (both the number of linked active vOTUs and increase in vOTU abundance), supporting the ‘kill-the-winner’ theory described in aquatic systems (Fig. 3; Fig. 4) [85]. In this Lotka–Volterra predator-prey framework, the most abundant host would become infected by viruses, leading to a decline in host abundance and an increase in the prevalence of their viruses, at least until microbial hosts with genetic resistance to the currently dominant viruses had time to respond. The dominance of Firmicutes hosts makes sense, since this diverse bacterial phylum is known for adaptations to anoxic conditions, including fermentation and creating endospores [86]. In our samples, all active infected Firmicutes contained fermentation genes for the capacity to produce ethanol, lactate, or both (Table S8). Bacteroidota were also an active and frequently infected bacterial phylum that increased in abundance through time; of this group, all the infected MAGs shared taxonomic affiliation to the order Bacteroidales. Most of these bacteria are obligate anaerobes and known for their diverse arrays of carbohydrate-active enzymes arranged into polysaccharide utilization loci and fermentation [87, 88]. Many active MAGs within this group had the genomic capacity for polysaccharide degradation (e.g., ICE 1 encoded pectin degradation protein KdgF) and all encoded genes for lactate fermentation (Table S8). After one year of subzero and anoxic incubation, Bacteroidales had become the most abundant active lineage in our soils and were infected by the most abundant active vOTUs (#s 23, 37, 41, 124, and 190), further supporting ‘kill-the-winner’ theory and highlighting viruses influencing host winter activities. While the abundance of active Bacteroidota and Firmicutes populations were increasing, C mineralization to CO_2_ occurred steadily throughout the incubation and vOTUs linked to these populations carried several AMGs that would support central carbon metabolism (Table S5). Together, these results suggest that these two bacterial lineages play an important role in over-winter C loss from these Artic peat soils, and the viruses that infect them likely shape both their population dynamics and functional impact.

Another way viruses can impact soil biogeochemistry is by infecting hosts that occupy different metabolic niche spaces. The Patescibacteria we identified (part of the Candidate phyla radiation; CPR) are known for their ultrasmall cell size with reduced genomes and most have a symbiotic relationship with Bacteroidota [89], suggesting direct interactions between these two lineages in our cryoecosystem. Patescibacteria are thought to resist phage infection by lacking or reducing the number of potential phage receptors on their cell membrane [90], but in our study, it was notably one of the most infected lineages and had none of the previously reported phage receptors (Table S8). The prevalence of these infections may be the result of their interactions with Bacteroidota. One member of the Patescibacteria was linked to an unlabeled vOTU with a CRISPR spacer match, suggesting this adapted immune system element was successful at protecting the host from infection. Typical CRISPRs are rarely found in CPR bacteria and recent work suggests this may be due to this group using a compact CRISPR-CasY system that is highly divergent to typical CRISPR systems [91]. Another infected and active bacterial phylum we observed was Proteobacteria, with only one active MAG from the class Micavibrionales. Little is known about this clade, and even less about their viruses, because they have an epibiotic lifestyle where they feed on other organisms to survive, making them difficult to culture. These predatory bacteria may be active under winter conditions by either consuming non-active or dead cells, or they may benefit from attaching to a host that can utilize alternative energy sources and recalcitrant organic matter [92]. We identified multiple active vOTUs infecting this lineage, revealing another indirect way that viruses may impact nutrient cycling and microbial diversity, via preying upon bacterial predators.

### Challenges associated with identifying virus ‘activity’

Heavy-water SIP has proven to be a robust method for identifying metabolically active microbes in soils [32, 33, 36]. In many ways, the approach is superior to other techniques that track activity such as RNA-to-DNA ratios, rRNA levels, bioorthogonal non-nanonical amino acid tagging (BONCAT) or other stable isotopes (e.g., ^13^C) because active microbes synthesize DNA when their cells divide, incorporating ^18^O, and therefore the DNA of all active microbes is labeled. Data from RNA studies can be hard to interpret as RNA levels often do not correlate with growth and often have weak or no correlation with proteins levels [93–95]. Compared to other isotopes as tracers, ^18^O labeling via heavy-water SIP does not rely on the microbe’s carbon use efficiency or prior knowledge of the microbe’s substrate preference [37].

The application of isotope tracers for direct assessment of activity is not as straightforward for viruses as it is for other microorganisms, and worthy of reasoned consideration. Characterizing activity for viruses is quite different compared to ‘free-living’ organisms because of their different infection cycles, their lack of metabolism, and the many states in which they can be present [19]. One of the main reasons to identify active entities is to help quantify their impact on their hosts and their environment, with the assumption that inactive entities have less of an impact but are still important [96]. For a microbe, this may be true, but for a virus there is a range in magnitude of impact for different host metabolic states that depends heavily on the infection cycle. Assessing the prevalence of an infection cycle, however, is challenging due to difficulties in quantifying lysogeny or virion abundance, burst size, diversity, and ecology [51].

Arctic soils, that predominantly exist under anoxic and subzero temperature conditions, might be generally considered a harsh environment, and would limit microbial growth. For this reason, we hypothesized most of our viruses would be temperate [51, 97] and they would be detected as proviruses. We did also see a high rate of lysogeny (43%), and in support of our hypotheses, the majority (80%) of temperate viruses were present as proviruses. Further, half of the active vOTUs linked to active MAGs (i.e., bacterial phyla Bacteroidota, Firmicutes, Patescibacteria, and Proteobacteria) were temperate viruses. The number of temperate viruses present in this study was higher than previously reported for desert Antarctic soils (4–20%) and almost identical (44%) to whole soil assays from temperate wetlands soils [98]. We hypothesize the increased incidence of temperate viruses is linked to low host abundances and environmental conditions, therefore increasing the potential for sporadic viral infection.

Temperate viruses can undergo lytic infection, where activity is identified by progeny viruses, or lysogenic infection, where activity is difficult to assess. A temperate virus that is undergoing lysogenic infection (present as a provirus) during a heavy-water SIP incubation would become isotopically enriched and depend on its host’s division rate for abundance. Active proviruses undergoing lysogenic infection need to be distinguished from viruses undergoing lytic infection because the effect of proviruses on the host metabolism (and therefore ecosystem) will not be as pronounced. This is because proviruses do not shut down and redirect host metabolism for progeny production during the lysogenic cycle as compared to the lytic cycle. During lysogeny, viruses still impact host metabolism via host gene regulation and acquisition of new virulence factors, but it is primarily for maintaining the provirus in the host genome [99]. Proviruses may also be labeled, but not currently active if they are proviral remnants of a past infection [100]. These remnants have no negative impact on host metabolism (beyond occupying genome space) but may confer some advantage as a gene transfer agent [51] or by contributing virulence factors which can still impact host physiology and metabolism [99]. SIP-enabled metagenomics alone cannot unequivocally identify a virus’ state (e.g., in maintenance mode or remnant), making it currently difficult to fully assess viral activity. Combining SIP-metagenomics with other approaches—such as a SIP-virome or induction assays—may identify vOTUs that have undergone lytic infection, and therefore provide a more holistic view of vOTU dynamics.

## Conclusions

Winter carbon losses in Arctic peat soils are estimated to be significant and growing, but the mechanisms that drive these losses are poorly understood. Using an ^18^O-heavy-water incubation under subzero, anoxic conditions, we found that bacteria and their viruses are active over the long winter months in northern peatlands, and these active populations drive significant CO_2_ fluxes. Our approach, SIP-targeted metagenomics, allowed us to move beyond a general catalogue of the genetic repertoire of these soil communities, and expose the specific population-level dynamics and functional capacities of the activity viral community. Given the high abundance of unlabeled bacteria and their viruses, these bacterial and viral populations would have been traditionally described first and foremost, thus occluding the characterization of the active bacteria and viruses in these soils. Despite the subzero temperatures and lack of oxygen in these peat soils, many active hosts and active viruses (both temperate and lytic) appear to be engaged in a surprisingly high level of biotic interactions and biogeochemical processing.

## Acknowledgements

We would like to thank the Millard lab for curating a list of Genbank bacteriophage genomes used for taxonomic identification of our vOTUs. Thanks to Jack McFarland and Monica Haw (USGS) for help and support with the experimental setup and maintenance. Sample collection and processing was supported by a US Geological Survey Mendenhall Fellowship to S.J.B., the Bonanza Creek LTER Program, jointly funded by NSF (DEB 1026415) and the USDA Forest Service Pacific Northwest Research Station (PNW01-JV112619320-16), and support from the USGS Climate R&D Program and AK Climate Science Center. Analysis of results was supported by a Lawrence Livermore National Laboratory, Laboratory Directed Research & Development grant (18-ERD-041) to S.J.B. and by LLNL’s U.S. Department of Energy, Office of Biological and Environmental Research, Genomic Science Program LLNL ‘Microbes Persist’ Scientific Focus Area (#SCW1632). Work conducted at PNNL, operated for the DOE by Battelle Memorial Institute, was conducted under Contract DE-AC05-76RLO1830. Work conducted at LLNL was conducted under the auspices of the US Department of Energy under Contract DE-AC52-07NA27344.

