## Supplemental text and figures 1-4 for "Ecology of active viruses and their bacterial hosts in frozen Arctic peat soil revealed with H_2_^18^O stable isotope probing metagenomics"

*Supplementary Information*

*Placing vOTUs in the context of global viruses*

We constructed a genome-based network to taxonomically classify our 332 vOTUs in the context of previously characterized viral genomes with ICTV-approved taxonomy [1]. The network identified 413 viral clusters (VCs; approximately genus-level taxonomy), 347 of which contained only RefSeq viruses, and 66 VCs that contained at least one of our vOTUs (Fig. S2; Table S3). Analysis of these 66 VCs indicated that the majority (54) contained only vOTUs specific to our Bonanza Creek LTER samples (16 vOTUs). And additional 192 of our vOTU were singletons with no supported gene-sharing connection to any other viral genome. We investigated the 12 VCs that contained members from both RefSeq and our dataset for higher taxonomic affiliation. All contained viruses from the order Caudovirales identifying 16 vOTUs as dsDNA bacteriophage with 11 Siphoviridae, 2 Podoviridae (including 1 crAss-like virus), and 3 Myoviridae. The host taxonomy of the RefSeq viruses spanned 4 bacterial phyla —Proteobacteria, Actinobacteria, Firmicutes, Bacteroidetes— all of which are included in the 10 bacterial phyla predicted for these vOTUs and the host phylum predicted for vOTU 56 and vOTU 237 (VCs 15 and 61, respectively) were Bacteroidetes and Firmicutes, and were corroborated by the nucleotide similarity matches (Table S9; Table S10). We identified VCs that contained multiple vOTUs and at least one vOTU with a taxonomic affiliation from our host-linkage analysis that we could leverage. In total 34 VCs also had taxonomic assignment from nucleotide similarity and three also with matching support from CRISPR spacer similarity. Of these, 16 previously unclassified vOTUs could be taxonomically classified and 7 VCs had multiple hosts linked to them. In all we were able to taxonomically classify 205 (62%) vOTUs using robust vOTU genomes and network analytics.

To improve the taxonomic affiliation of our vOTUs, we created a second network that incorporated 5-fold more viral genomes, adding genomes drawn from environmental surveys where the genome completeness is unknown. Previous work suggests that more high-quality viral genomes can help to connect singletons and improve connections to new or existing VCs [1]. We drew upon thousands of near-complete viral genomes in Genbank (May 30, 2020) [2] that were manually curated to meet the necessary robust criteria (i.e., validated to be viral origin, ≥10kb, and dereplicated) to be included in a network. In this second network, we identified 1 736 VCs, 60 of which contained at least one vOTU, 55 only contained vOTUs (Fig. S4; Table S11), and 202 singletons. Notably, adding more viral genomes to the network did increase the number of VCs, but decreased the number of VCs that contained a vOTU reported here and resulted in only 130 vOTUs in the network. A comparison of the two networks revealed 123 shared vOTUs and 24 unique (17 only clustered in the first network and 7 in the second network). The second, larger network provided taxonomic affiliation to four of the vOTUs, including new assignments for vOTUs 36 and 122 (both are Caudovirales, Myoviridae; potential host phylum Bacteroidetes), the same taxonomic and host assignments for vOTUs 93, 146, and 228, and the same viral taxonomic assignment for vOTU 168, but expanded the host phylum assignment from Actinobacteria (as reported in network 1), to also include Firmicutes.

### Supplementary Methods

#### *Generation of gene-sharing networks*

A gene-sharing network was generated to compare the vOTUs globally to validated viral genomes following the suggested applications [3] on CyVerse [4] and documentation on Protocols.io (https://www.protocols.io/view/applying-vcontact-to-viral-sequences-and-visualizi-x5xfq7n). Briefly, a gene-to-contig file was created using the vContact2-Gene2Genome v1.1.0 (default parameters) application and a Prodigal amino acid protein file as input source type. To cluster vOTUs, we used vConTACT2 v0.9.8 [1] with default parameters except for: i) < 0.8 overlap for viral clusters (VCs); ii) multi merge method for highly overlapping VCs, and iii) NCBI Bacterial and Archaeal Viral RefSeq v85 with only ICTV (International Committee on Taxonomy of Viruses)-approved taxonomy as a reference database. The network was visualized with Cytoscape v3.7.2 [5], using an edge-weighted spring embedded model and removing duplicated nodes.

A second more inclusive network was generated using a viral database downloaded from the Millard lab website (http://millardlab.org/) along with a mapping file for coloring VCs and uploaded to CyVerse for processing (using the same applications and parameters listed above). The viral database consisted of more than 9,000 robust (≥10kb) or complete bacteriophage genomes, extracted from Genbank on May 31, 2018.

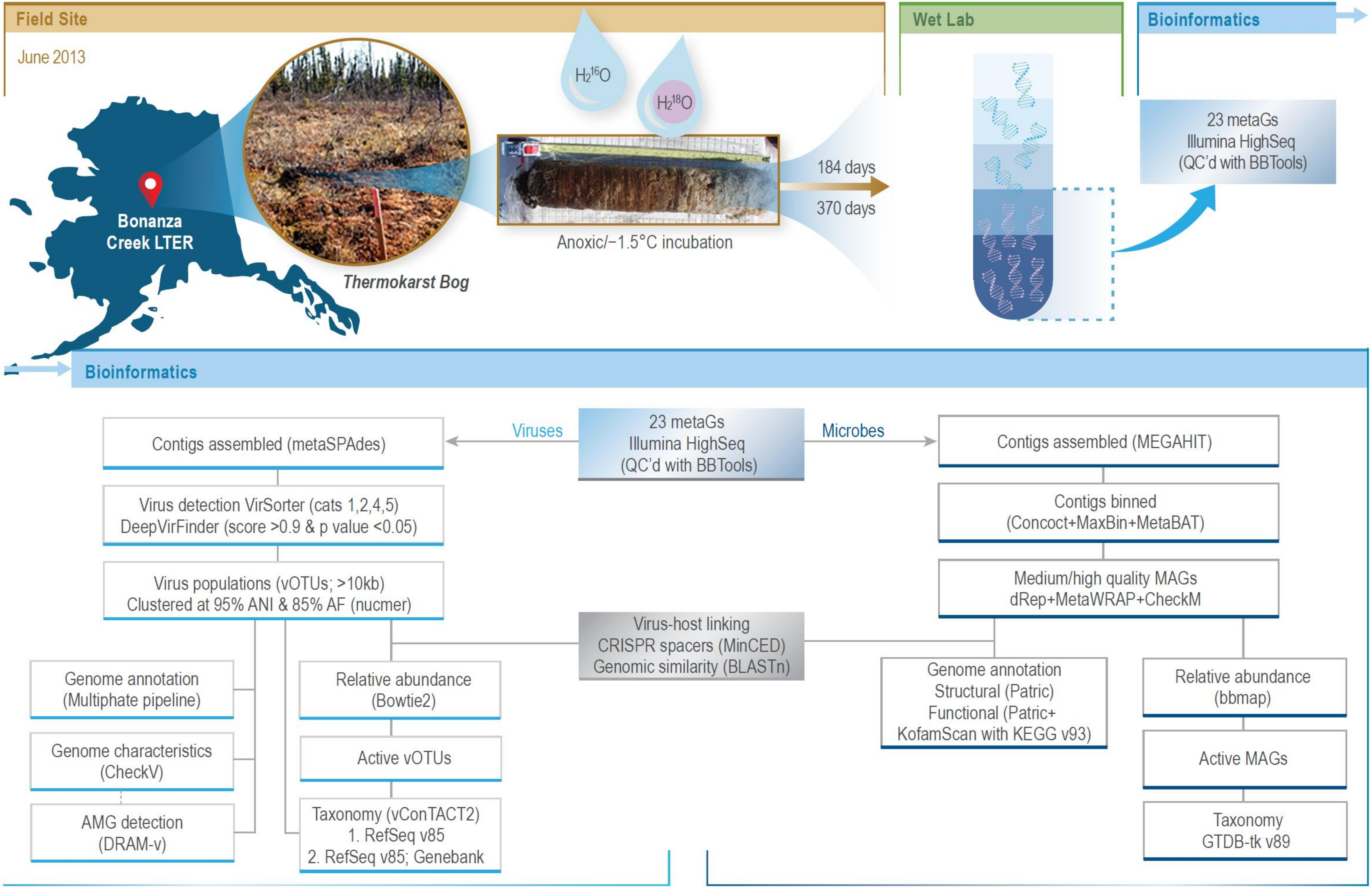
Field Site
June 2013
Bonanza
Creek LTER
Thermokarst Bog
$H_2{}^{16}O$
$H_2{}^{18}O$
Anoxic/−1.5°C incubation
184 days
370 days
Wet Lab
Bioinformatics
23 metaGs
Illumina HighSeq
(QC'd with BBTools)
Bioinformatics
Viruses
Microbes
Contigs assembled (metaSPAdes)
Virus detection VirSorter (cats 1,2,4,5)
DeepVirFinder (score >0.9 & p value <0.05)
Virus populations (vOTUs; >10kb)
Clustered at 95% ANI & 85% AF (nucmer)
Genome annotation
(Multiphate pipeline)
Genome characteristics
(CheckV)
AMG detection
(DRAM-v)
Relative abundance
(Bowtie2)
Active vOTUs
Taxonomy (vConTACT2)
1. RefSeq v85
2. RefSeq v85; Genebank
Virus-host linking
CRISPR spacers (MinCED)
Genomic similarity (BLASTn)
Contigs assembled (MEGAHIT)
Contigs binned
(Concoct+MaxBin+MetaBAT)
Medium/high quality MAGs
dRep+MetaWRAP+CheckM
Genome annotation
Structural (Patric)
Functional (Patric+
KofamScan with KEGG v93)
Relative abundance
(bbmap)
Active MAGs
Taxonomy
GTDB-tk v89

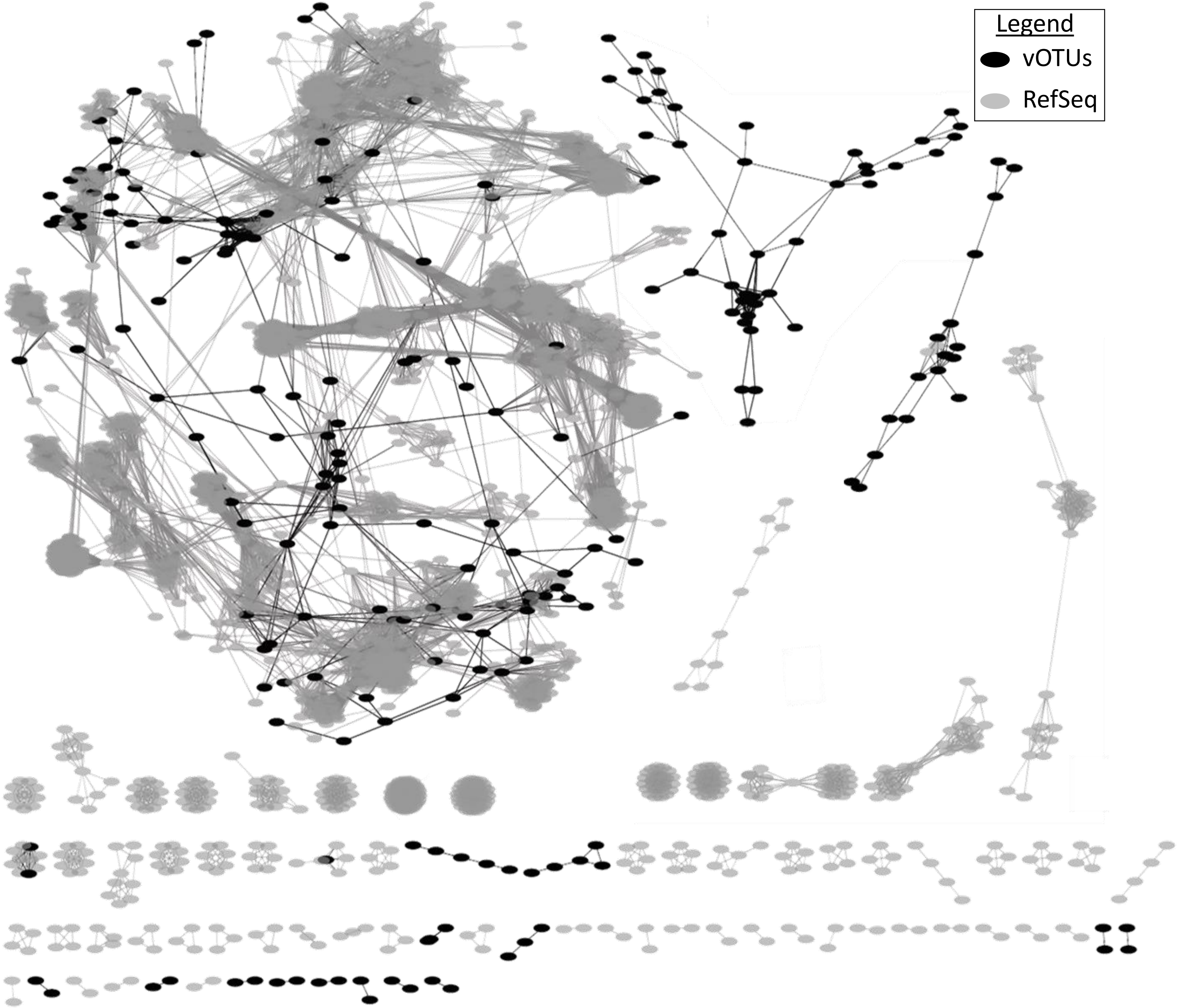

Legend
vOTUs
RefSeq

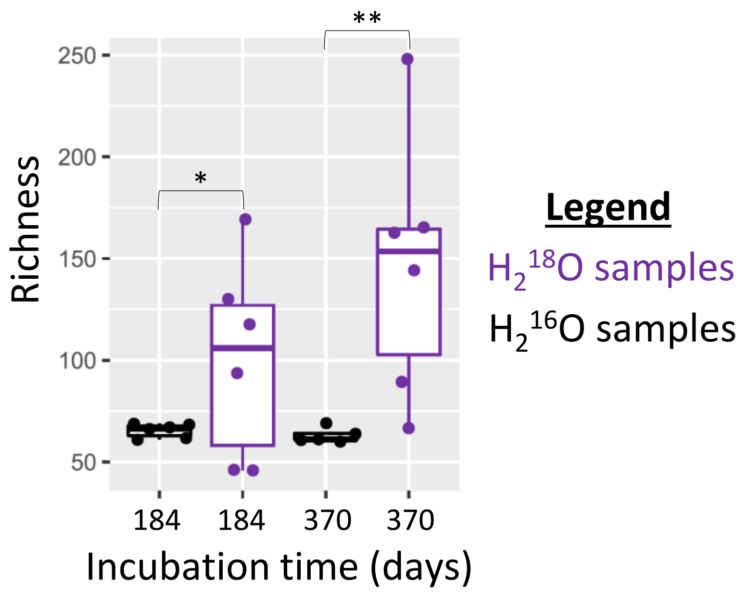

Richness
250
200
150
100
50
*
**
184
184
370
370
Incubation time (days)
Legend
$H_2^{18}O$ samples
$H_2^{16}O$ samples

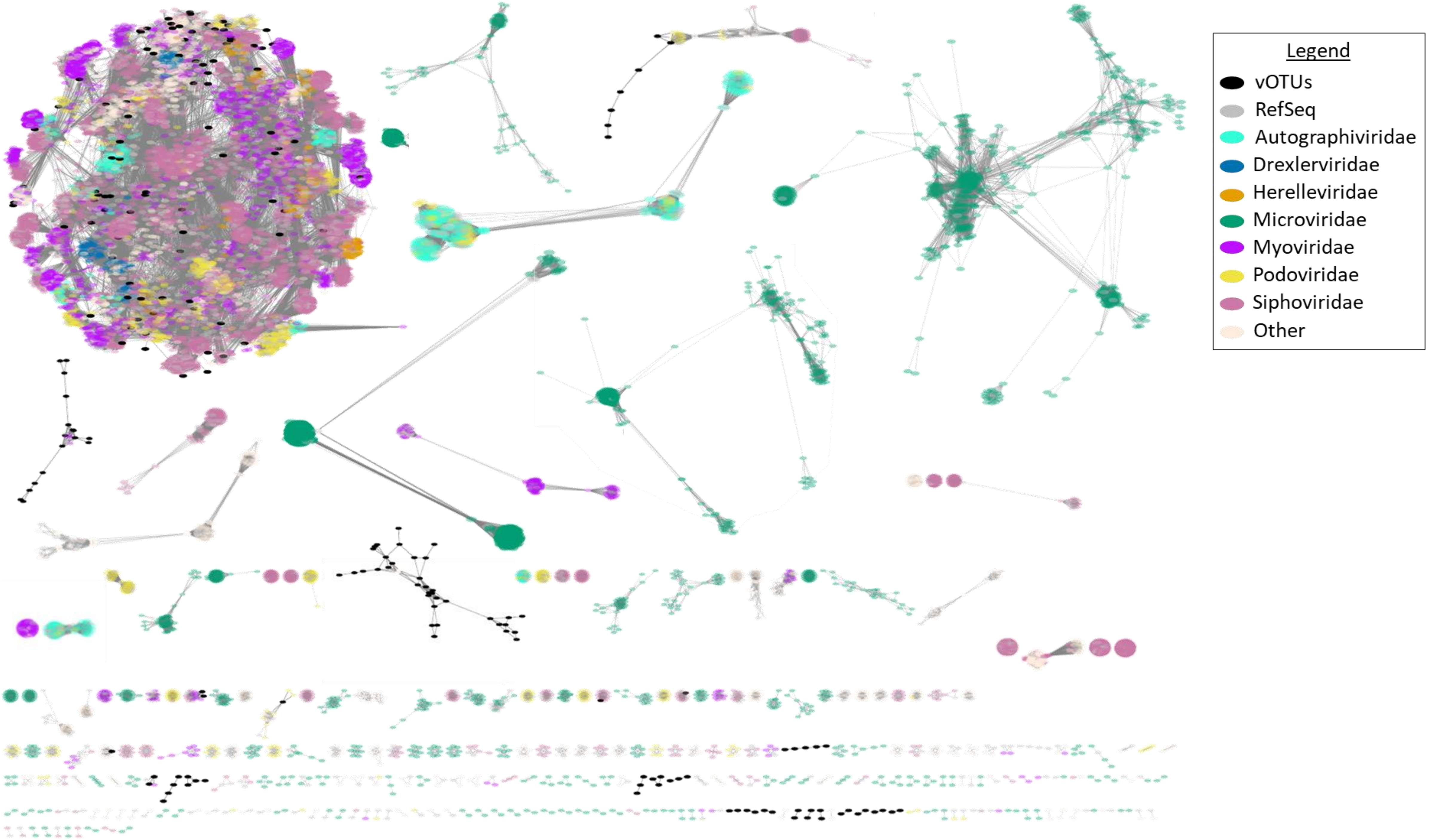

Legend
vOTUs
RefSeq
Autographiviridae
Drexlerviridae
Herelleviridae
Microviridae
Myoviridae
Podoviridae
Siphoviridae
Other
